# ADAR2-Mediated RNA Editing Promotes TDP-43 Nuclear Export and Alters RNA Binding

**DOI:** 10.64898/2026.06.22.730622

**Authors:** Stephen Moore, Dominic L. Julian, Eric Alsop, Lauren M. Gittings, Ileana Lorenzini, Michael McMillan, Sam Macklin-Isquierdo, Erik Lehmkuhl, Petr Kalab, Danielle de Paula Moreira, Lindsey Hayes, Christopher Donnelly, Sami J. Barmada, Daniela C. Zarnescu, Kendall Van Keuren-Jensen, Rita Sattler

## Abstract

**BACKGROUND:** TAR DNA binding protein - 43 (TDP-43) nuclear loss is a pathological hallmark of amyotrophic lateral sclerosis (ALS), frontotemporal dementia (FTD), and related neurodegenerative disorders. While the consequences of TDP-43 dysfunction have been well-characterized, the mechanisms driving TDP-43 mislocalization remain poorly understood. Previous observations of altered localization and function of the adenosine-to-inosine (A-to-I) RNA editing enzyme adenosine deaminase acting on RNA 2 (ADAR2) in ALS/FTD tissue prompted us to investigate whether dysregulated RNA editing contributes to pathological TDP-43 nucleocytoplasmic trafficking.

**METHODS:** TDP-43 cytoplasmic mislocalization was assessed following ADAR2 and TDP-43 co-overexpression in HEK293T cells and a *Drosophila* model co-overexpressing human TDP-43 and *dADAR* in motor neurons. We further evaluated TDP-43 mislocalization through both HeLa cell assays and interspecies heterokaryon assays. Next, we assessed TDP-43 binding to A-to-I edited RNA oligomers through electrophoretic mobility shift assays (EMSAs), and investigated inosine-containing RNAs *in vivo* via TDP-43 RNA immunoprecipitation followed by sequencing (RIP-seq) datasets from human TDP-43-expressing *Drosophila*. Finally, RNAseq and enhanced cross-linking and immunoprecipitation (eCLIP-seq) were performed in SH-SY5Y cells overexpressing three ADAR2 variants with differing editing activity to identify editing-related transcriptional alterations and RNAs differentially bound to TDP-43.

**RESULTS:** ADAR2 overexpression reduced the nucleocytoplasmic (N:C) ratio of TDP-43 in HEK293T cells in a ADAR2 catalytic activity- and TDP-43 RNA-binding capacity-dependent manner. *Drosophila* motor neurons overexpressing *dADAR* also exhibited decreased nuclear TDP-43. Interspecies heterokaryons and permeabilized HeLa cell assays demonstrated that catalytically active ADAR2 and synthetic inosine-containing RNA oligomers, respectively, enhance nuclear export of endogenous TDP-43. EMSAs revealed preferential binding of TDP-43 to inosine-containing RNAs relative to unedited RNAs, and analysis of *Drosophila* RIP-seq datasets demonstrated enrichment of edited transcripts within TDP-43-bound RNAs. Finally, RNAseq and eCLIP-seq analyses identified editing-dependent alterations in gene expression and TDP-43 RNA-binding profiles in SH-SY5Y cells overexpressing active ADAR2 variants.

**CONCLUSIONS:** Together, our findings identify A-to-I RNA editing as a previously unrecognized regulator of TDP-43 localization and RNA interactions. These results support a model where altered RNA editing modifies TDP-43-RNA interactions, promoting increased nuclear export of TDP-43. Broadly, our work highlights RNA editing dysregulation as a potential contributor to early pathogenic mechanisms underlying TDP-43 proteinopathies.

## BACKGROUND

Transactive response DNA Binding Protein of 43 kilodaltons (TDP-43) is a ubiquitously expressed RNA binding protein that performs essential roles in RNA metabolism, including RNA splicing, trafficking, and stability (1–4). Nuclear depletion and cytoplasmic accumulation of TDP-43 constitute the principal neuropathological hallmark in 97% of amyotrophic lateral sclerosis (ALS) and half of all frontotemporal dementia (FTD) patients (5–7). TDP-43 mislocalization has also been reported in multiple neurodegenerative disorders including Alzheimer’s disease, limbic-predominat age-related TDP-43 encephalopathy (LATE), Parkinson’s disease, and Huntington’s disease, underscoring the broad clinical significance of investigating TDP-43 dysfunction (8–11). Extensive work has defined the downstream consequences of TDP-43 dysfunction, particularly its effects on RNA processing. Loss of nuclear TDP-43 functions promotes the use of cryptic splice or polyadenylation sites resulting in altered RNA expression and production of cryptic protein products (12–17,17–19). In contrast, the upstream molecular events that initiate TDP-43 nuclear depletion and cytoplasmic accumulation remain poorly understood. Identification of these initiating mechanisms could reveal new therapeutic opportunities to prevent or delay TDP-43 proteinopathy.

Previous work from our laboratory demonstrated that a GGGGCC hexanucleotide repeat expansion in the *C9ORF72* (C9) gene, the most common genetic form of ALS and FTD (C9 ALS/FTD) (20,21) leads to the cytoplasmic accumulation of the RNA editing enzyme adenosine deaminase acting on RNA 2 (ADAR2) and an altered RNA editing profile in ALS/FTD brain and spinal cord tissue (22). ADAR2, along with ADAR1, can bind to double-stranded regions of RNAs and catalyze the deamination of adenosine nucleotides to inosines, which alters RNA secondary structure and function; this occurs in conjunction with the central nervous system-specific and catalytically inactive ADAR3, which exerts a regulatory role on its other family members (23–26). Notably, C9 ALS/FTD human spinal cord motor neurons exhibiting cytoplasmic ADAR2 accumulation frequently also contained cytoplasmic TDP-43 inclusions, suggesting a potential pathological relationship between these two RNA-regulating proteins in neurodegeneration.

Here, we investigated whether aberrant A-to-I RNA editing acts as an upstream trigger of pathologic TDP-43 nucleocytoplasmic trafficking. Indeed, we have identified that ADAR2 mediated A-to-I RNA editing activity disrupts normal TDP-43 nucleocytoplasmic shuttling and leads to TDP-43 cytoplasmic aggregation. In cultured cells, co-expression of ADAR2 and TDP-43 induced cytoplasmic TDP-43 accumulations, while deletion of the ADAR2 catalytic domain or disruption of TDP-43 RNA binding prevented this mislocalization. In *Drosophila*, co-expression of a *Drosophila* A-to-I editing enzyme (dADAR) and human TDP-43 similarly promoted TDP-43 cytoplasmic accumulation in motor neurons. Using two orthogonal nuclear export assays we further demonstrated that the rate of endogenous TDP-43 nuclear export is increased either by ADAR2 overexpression or by exposure to synthetic A-to-I edited RNA in the absence of ADAR2 overexpression. We also highlight the ability of TDP-43 to bind to inosine-containing RNA oligomers *in vitro* utilizing electrophoretic mobility shift assays (EMSAs), and further confirm this phenomenon *in vivo* through a bioinformatic analysis of TDP-43 RNA immunoprecipitation followed by sequencing (RIP-seq) data from *Drosophila* expressing humanized TDP-43. Finally, we utilized bulk RNA sequencing (RNAseq) and enhanced cross-linking and immunoprecipitation followed by RNA sequencing (eCLIP-seq) to reveal novel editing-dependent transcriptional alterations and TDP-43 binding sites in SH-SY5Y cells overexpressing ADAR2 variants with differing RNA editing capabilities.

Together, these findings demonstrate that A-to-I RNA editing is a novel upstream trigger of nuclear TDP-43 export and ultimately TDP-43 dysfunction. More broadly, they suggest that altered RNA editing may contribute to TDP-43 proteinopathy across multiple neurodegenerative diseases and highlight RNA-editing pathways as potential therapeutic targets.

## METHODS

### TDP-43 and ADAR2 Overexpression in HEK293T

HEK293T cells were cultured in DMEM with 10% FBS and 1% Pen-Strep at 37°C, 5% CO₂. Cells were seeded on poly-L-lysine–coated coverslips (1×10_ cells/well, 24-well plate) and transfected at ∼70% confluency with 0.5 µg plasmid using PEI (3:1 PEI:DNA) in OptiMEM. Media was changed after 5 h, and cells were fixed with 4% PFA the next day for imaging.

### Cloning of ADAR2ΔDA-mCherry and NLS-mCherry

ADAR2 constructs were generated from a GeneCopoeia plasmid (ADAR2-PEZ-LV111) using restriction digestion (EcoRI, BamHI, ClaI) and NEB HiFi assembly. PCR-amplified fragments were gel purified and assembled per manufacturer instructions. Constructs were transformed into Stbl3 E. coli, selected on ampicillin plates, expanded in LB broth, and verified by restriction digestion.

### Quantification of TDP-43 nuclear:cytoplasmic ratio

Confocal images (Zeiss LSM800, 63X objective) were acquired using identical settings. Maximum intensity projections were analyzed in Zen software. Nuclear ROIs were defined by DAPI, and cytoplasmic ROIs (3 per cell) by WGA staining. Mean fluorescence intensities were used to calculate nuclear:cytoplasmic (N:C) ratios (log₂ transformed). Statistical analysis was performed using one-way ANOVA (GraphPad Prism).

### ADAR2 Lentivirus Generation

ADAR2 constructs (FL-ADAR2; ΔNLS-ADAR2; Tet-On-ADAR2 WT; Tet-On-ADAR2-E396A; Tet-On-ADAR2-E488Q) were transformed into Stbl3 cells, purified, and transfected into HEK293T cells (15 cm dishes, 10×10_ cells) using a three-plasmid lentiviral system (2:4:1 transfer:envelope:packaging). Media was changed after 5 h, and virus was collected at 24, 48, and 72 h and concentrated using Lenti-X Concentrator (Takara).

### RNA immunoprecipitations in *Drosophila*

*Drosophila* RNA immunoprecipitations (RIP) and RNA isolation were previously described (27). Briefly, 100 third instar larvae overexpressing human TDP-43-YFP were collected and prepared for immunoprecipitation. Anti-GFP antibodies were used to immunoprecipitate TDP-43, and RNA was isolated and sequenced from both input and IP fractions. *Drosophila*-specific RNA editing sites were downloaded from the REDI portal (28). RNA editing ratios were then quantified at each editing site as previously described (22).

### Overexpression of TDP-43 and ADAR in *Drosophila*

*Drosophila* lines D42-GAL4 (BDSC_8816) (29), UAS-hTDP-43^WT^-YFP (30), and UAS-hADAR/CyO Kr::GFP (31) were used to express the human TDP-43-YFP and *Drosophila* ADAR (30–32). All *Drosophila* stocks were kept on standard yeast/cornmeal/molasses food and incubated at 25°C. The GAL4-UAS crosses were kept at 22°C with a 12h:12h light:dark regime.

### Image analysis of *Drosophila* ventral nerve cord neurons

Ventral nerve cords (VNCs) from third instar larvae expressing TDP-43 alone or with ADAR were dissected in 1× PBS, fixed in 4% PFA, permeabilized in 0.3% PBST, and blocked in 1% BSA/2% normal goat serum. TDP-43-YFP was detected using anti-GFP-FITC, and nuclei were stained with Hoechst. Images were acquired on a Leica SP8 confocal microscope (63× oil objective) with identical settings and 1 µm z-stacks. For analysis, a single optical section per VNC containing ≥5 motor neurons was selected. Nuclear ROIs were defined by Hoechst staining, and whole-cell ROIs by GFP signal. Mean fluorescence intensities were quantified in ImageJ (v1.54p), and nuclear-to-whole-cell TDP-43 ratios were calculated for each neuron. Outliers were removed using the ROUT test (Q = 0.5%), and statistical comparisons were performed using Welch’s t-test in GraphPad Prism (v11).

### Interspecies Heterokaryon Nuclear Export Assay

Nuclear export was assessed using an interspecies heterokaryon assay as previously described with minor modifications (33–35). SH-SY5Y cells expressing endogenous TDP-43-eGFP (36) were transduced with ADAR2-mCherry or ADAR2ΔDA-mCherry, then mixed 1:1 with mouse embryonic fibroblasts (MEFs) and seeded on fibronectin-coated coverslips. Cells were treated with cycloheximide (100 µg/mL, 2 h) to inhibit protein synthesis and fused by incubation with 50% polyethylene glycol (PEG) for 2 min, followed by washing in PBS containing cycloheximide. Heterokaryons were incubated in complete media with cycloheximide for defined time points to allow nucleocytoplasmic shuttling, then fixed with 4% PFA and processed for immunocytochemistry as described previously(22). Cells containing one human and one mouse nucleus were identified based on Hoechst staining and species-specific nuclear morphology; multinucleated cells were excluded. TDP-43-eGFP nuclear export was quantified by measuring GFP intensity in donor and recipient nuclei using Cell Profiler (37).

### Electromobility shift assays

Electromobility shift assays (EMSAs) were performed as previously described (38,39). Briefly, recombinant His-SUMO-tagged WT TDP-43 was isolated from *E. coli* as in (38), then mixed with 100 pM labeled oligonucleotide (IDT) in binding buffer (12.5 mM HEPES, pH 7.8, 50 mM KCl, 2.5 mM MgCl2, 0.5 mM TCEP, 25 μg/mL BSA, 0.01% NP-40) supplemented with 50% glycerol and 1μg/μl poly-dIdC, for 5 min at 4°C, followed by incubation at room temperature for 20 min. Reactions were then subjected to electrophoresis on 6% acrylamide gels and were imaged using the LI-COR Odyssey platform. Relative intensities for bound and unbound probe were determined in Fiji, and dissociation constants (Kd) and Hill coefficients were calculated using Prism (GraphPad) as described in (39).

### Nuclear export assays in HeLa cells

HeLa cell nuclear export assays were performed as previously described (40). Briefly, HeLa cells were cultured on coverslips, permeabilized with Digitonin and then incubated with various RNA oligomers for 30 minutes at 25°C. Following incubation, cells were fixed and stained for TDP-43 and the nuclear intensity of TDP-43 signal was quantified and compared for each RNA oligomer at increasing concentrations.

### Generation of stable ADARB1 lentivirus-transduced SH-SY5Y cells

SH-SY5Y neuroblastoma cells were cultured in 1X DMEM + 10% FBS and plated at 1.0×10^6^ cells in a 10 cm^2^ cell culture treated dish, and were transduced with Tet-On-ADARB1 lentiviral (ADARB1-LV) constructs (ADAR2^WT^, ADAR2^E396A^, ADAR2^E488Q^; GeneCopoeia) at an MOI:10. Lentivirus-containing media was discarded 24 hours post-transduction, and was replaced with media containing 0.5 mg/mL of Geneticin and media changes were performed every other day for selection of transduced cultures for the next 7 days.

### eCLIP Methods

Stable ADARB1-LV-transduced SH-SY5Y cells were grown on 15 cm^2^ dishes and cultured until ∼75% confluency. Cultures were either left alone or 2 µg/mL Doxycycline (ThermoFisher, #J67043.AD) was added to the cultures for 48h to activate lentiviral overexpression. Cells were cross-linked on ice at 6000 J/cm^2^ using 254 nm UV light, and immediately collected, spun down, and flash-frozen on dry ice. Samples were processed utilizing the previously published single-end seCLIP protocol with modifications (41). Pre-validated TDP-43 antibodies (Proteintech, #10782-2-AP) were used for immunoprecipitation.

### eCLIP Data Processing

Raw sequence files had unique molecular identifiers (UMIs) on the eCLIP cDNA adapters pruned from read sequencing using umi_tools (v0.5.1) (42,43). 3’-adapters were trimmed from reads using cutadapt (v2.7), and reads shorter than 18bp in length were removed (44). Reads were mapped to a database of human repetitive elements and rRNA sequences compiled from Dfam (45) and Genbank (46). All non-repeat mapped reads were mapped to the human genome (hg38) using STAR (v2.6.0c) (47). PCR duplicates were removed using umi_tools (v0.5.1) by utilizing UMI sequences from the read names and mapping positions.

### RNA Sequencing Methods

Total RNA was isolated from frozen cell pellets using RNeasy Plus Mini kit (Qiagen). Cell pellets were resuspended in Buffer RLT Plus supplemented with BME. Cells were homogenized using Qiashredder columns and the remaining of the protocol was following using the manufacturer’s suggested protocol for total RNA. RNA was quantified using Ribogreen (ThermoFisher) and RNA integrity was determined using high sensitivity RNA assay on the Tapestation 4200 (Agilent).

### Whole Transcriptome Library Preparation and Sequencing

100ng of total RNA was used to generate libraries for sequencing using the Takara SMART-Seq Total RNA High Input (RiboGone Mammalian). Library preparation was performed following manufacturer’s recommended guidelines. 13 cycles of PCR was used to amplify libraries. Libraries were quantified using high sensitivity d1000 assay on the Tapestation 4200 (Agilent). 1μL of each library was used in a pool sequenced on the iSeq100. Equimolar pools were generated based on the reads from the iSeq100. The pool was quantified using the high sensitivity d1000 assay (Agilent) and qPCR (Roche). Libraries were sequenced on the NovaSeqX Plus with instrument read settings at 100 cycles for read 1 and 2 and 8 cycles for the index cycles with a read depth of at least 50M paired end reads.

### RNA Sequencing Data Analysis

Fastq files were generated from the raw sequence files using bcl2fastq (Illumina) using default parameters. Reads were trimmed with cutadapt v1.17 according to kit recommendations. Trimmed fastq files were then aligned to the GRCh38 genome with STAR v2.6.1d with the following parameters: --runMode alignReads --outSAMtype BAM Unsorted --outSAMmode Full -- outSAMstrandField intronMotif --outFilterType BySJout --outSAMunmapped Within -- outSAMmapqUnique 255 --outFilterMultimapNmax 20 --outFilterMismatchNmax 999 -- outFilterMismatchNoverLmax 0.1 --alignMatesGapMax 1000000 --seedSearchStartLmax 50 -- alignIntronMin 20 --alignIntronMax 1000000 --alignSJoverhangMin 18 --alignSJDBoverhangMin 18 -- chimSegmentMin 18 --chimJunctionOverhangMin 18 --outSJfilterOverhangMin 18 18 18 18 --alignTranscriptsPerReadNmax 50000. Following genome alignment, reads were counted with featureCounts v1.6.3, (part of the subread package) using a non-redundant genome annotation combined from GENCODE 47 and LncBook v2,1 and the following parameters: -p -t exon -g gene_id. Additionally, the strandedness parameter was passed to featureCounts. Raw gene count table for use with downstream analyses were generated using a python script that combined all featureCounts output files into a single TSV file. Normalized bulk RNA-sequencing counts were filtered to retain protein-coding genes, and the 10,000 most variable protein-coding genes across samples were selected for downstream analysis. Principal component analysis (PCA) was performed using the prcomp function in R.

### RNA Editing Analyses

A-to-I RNA editing locations were determined by downloading the complete list of human editing locations from the REDiportal database (∼16M known editing locations). Samtools mpileup was run on each BAM file against the REDiportal list of editing locations to determine read coverage at each editing site as well as an editing ratio (number of reads edited A-to-I divided by total number of reads at the genomic location). RNA editing sites were subjected to quality-control filtering to remove low-information sites with minimal editing or little variation across samples and only editing calculations for sites that had an average coverage of at least 10 reads were included. This resulted in 82,437 editing sites incorporated in downstream analyses. To prioritize RNA editing sites, editing frequencies were analyzed using weighted gene co-expression network analysis (WGCNA). Modules exhibiting coordinated changes in editing following ADAR2 induction were identified, and editing sites belonging to modules associated with increased editing in WT and E488Q cells relative to E396A cells were selected for subsequent eCLIP analyses.

Strand-matched eCLIP signal was extracted within a 500-nt window centered on each editing site (±250 nt). TDP-43 occupancy was quantified as the difference between mean signal within a central 50-nt region surrounding the editing site (−25 to +25 nt) and the mean signal in flanking background regions (−250 to −100 nt and +100 to +250 nt). To identify candidate editing-dependent TDP-43 binding events, an editing-dependent occupancy score was calculated for each editing site. Editing site were ranked according to an editing-dependent TDP-43 occupancy score defined as ΔWT + ΔE488Q – ΔE396A, where Δ indicated the doxycycline induced change in normalized TDP-43 eCLIP signal.

### Writing Assistance within Manuscript

The manuscript has been written entirely by the authors, with the utilization of artificial intelligence-derived large language models (LLMs) such as ChatGPT-5.5 for assistance with abbreviating certain sections of the manuscript. No original portion of this manuscript originated from the utilization of ChatGPT or any other LLM.

## RESULTS

### ADAR2 induces TDP-43 mislocalization via a previously unrecognized RNA editing-dependent mechanism

Following our previous observation that cytoplasmic ADAR2 accumulation is associated with TDP-43 pathology in human ALS/FTD tissue, we asked whether ADAR2-associated A-to-I RNA editing might directly influence TDP-43 localization. To test this, we co-expressed human TDP-43 containing an N-terminal GFP tag (TDP-43-GFP) together with human ADAR2 containing an N-terminal mCherry tag in HEK293T cells (Figure 1A). Unexpectedly, cells co-expressing ADAR2 and TDP-43 frequently exhibited cytoplasmic accumulation of TDP-43-eGFP, reminiscent of the pathological TDP-43 redistribution observed in ALS/FTD patient tissue(6,7). Quantification of the nuclear-to-cytoplasmic (N:C) ratio of TDP-43-eGFP 24 h post-transfection revealed a reduction in the N:C ratio in cells with positive GFP and mCherry signals, which is consistent with increased cytoplasmic accumulation of TDP-43 (Figure 1B,C; p < 0.0005).

**Figure 1.**
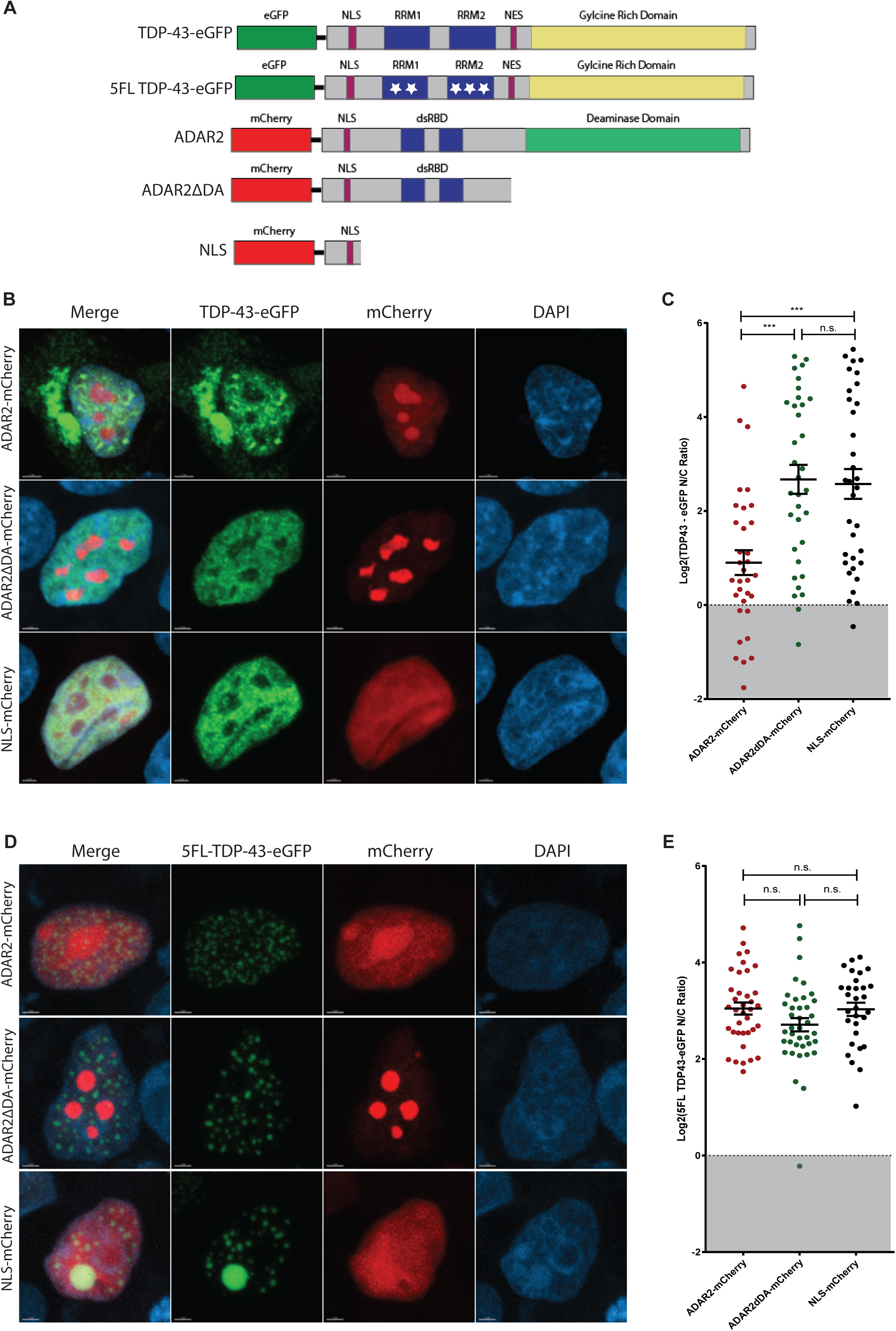
ADAR2 activity and functional TDP-43 RNA recognition motifs are required for editing-induced TDP-43 nuclear export **a)** Constructs used to overexpress TDP-43-eGFP, 5FL-TDP-43-eGFP ADAR2-mCherry, ADAR2ΔDA-mCherry, and NLS-mCherry in HEK293T cells. **b)** Representative images of TDP-43-eGFP mislocalization with ADAR2-mCherry overexpression but not with ADAR2ΔDA-mCherry or NLS-mCherry; scale bar = 2µm. **c)** Log2(TDP-43-eGFP N/C Ratio) is decreased in HEK293T cells overexpressing ADAR2-mCherry; n=30 cells/group, one-way ANOVA, ***p<0.0005. **d)** Representative images of 5FL-TDP-43-eGFP with overexpression of ADAR2-mCherry, ADAR2ΔDA-mCherry, and NLS-mCherry; scale bar = 2µm. **e)** No changes in the N/C ratio of 5FL-TDP-43-eGFP in HEK293T cells, irrespective of co-transfection; n=30 cells/group, one-way ANOVA, n.s. = not significant.

Because overexpression of TDP-43 alone can promote mislocalization, we wondered whether co-expression with another RNA-binding protein might nonspecifically enhance this effect, or whether ADAR2 RNA-editing activity directly influences TDP-43 nucleocytoplasmic trafficking. To distinguish between these possibilities, HEK293T cells were co-transfected with TDP-43-GFP and either a catalytically inactive ADAR2 construct lacking the deaminase domain (ADAR2ΔDA) or a control construct containing only the ADAR2 nuclear localization sequence (NLS-mCherry). Unlike full-length ADAR2, neither ADAR2ΔDA nor NLS-mCherry altered the nuclear-to-cytoplasmic ratio of TDP-43-GFP (Figure 1B,C; p > 0.05), indicating that ADAR2 catalytic activity, rather than nonspecific co-expression, is required to drive TDP-43 mislocalization. To further control for the possibility that complete deletion of the ADAR2 deaminase domain introduced unintended effects unrelated to editing activity, we generated two additional ADAR2-mCherry variants containing point mutations within the catalytic domain. ADAR2^E396A^ exhibits reduced editing activity (48), whereas ADAR2^E488Q^ displays enhanced editing activity (49). Co-expression of ADAR2^E396A^ with TDP-43-GFP did not alter the N:C ratio of TDP-43, whereas ADAR2^E488Q^ reduced the N:C ratio to a degree similar to wild-type ADAR2 (Supplementary Figure 1; p < 0.05)

Multiple studies have suggested that RNA binding by TDP-43 can influence its subcellular localization and nucleocytoplasmic trafficking (40,50). In addition, chemical modifications of RNA, such N6-methyladenosine, have been shown to alter TDP-43 RNA interactions and modulate its localization (10,51,52). These observations raised the possibility that A-to-I edited RNA may similarly influence TDP-43 behavior through altered RNA binding. To determine whether RNA binding by TDP-43 is required for ADAR2-dependent mislocalization, we utilized a TDP-43 construct containing five point mutations across both RNA recognition motifs (RRMs), which abolish RNA binding (5FL-TDP-43-GFP; Figure 1A) (50,53–55). Co-transfection of 5FL-TDP-43-GFP with either wild-type ADAR2, ADAR2ΔDA, or NLS-mCherry did not alter the N:C ratio of TDP-43 (Figure 1D,E; p > 0.05). These findings indicate that the ability of TDP-43 to bind RNA is required for ADAR2-dependent cytoplasmic redistribution.

### *dADAR* overexpression promotes nuclear loss of TDP-43 in *Drosophila* motor neurons

To investigate whether RNA editing similarly influenced TDP-43 localization in an *in vivo* neuronal context, we examined if increased ADAR activity could alter TDP-43 localization in *Drosophila* motor neurons. Ventral nerve cords (VNCs) were collected from larvae overexpressing human TDP-43 alone or co-expressing human TDP-43 together with *Drosophila* ADAR (*dADAR*), an orthologue of human ADAR2. Immunofluorescent analysis revealed marked cytoplasmic redistribution of TDP-43-YFP in flies co-expressing dADAR, whereas flies expressing human TDP-43-YFP alone retained predominantly nuclear TDP-43-YFP localization (Figure 2E,F). Quantification of the nuclear and whole cell TDP-43-YFP ratio confirmed a reduction in nuclear TDP-43 in the *dADAR* co-expression group relative to TDP-43-YFP alone (Figure 2G; ****p = 0.0001). These findings indicate that enhanced RNA-editing activity is sufficient to promote TDP-43 mislocalization in a model organism.

**Figure 2.**
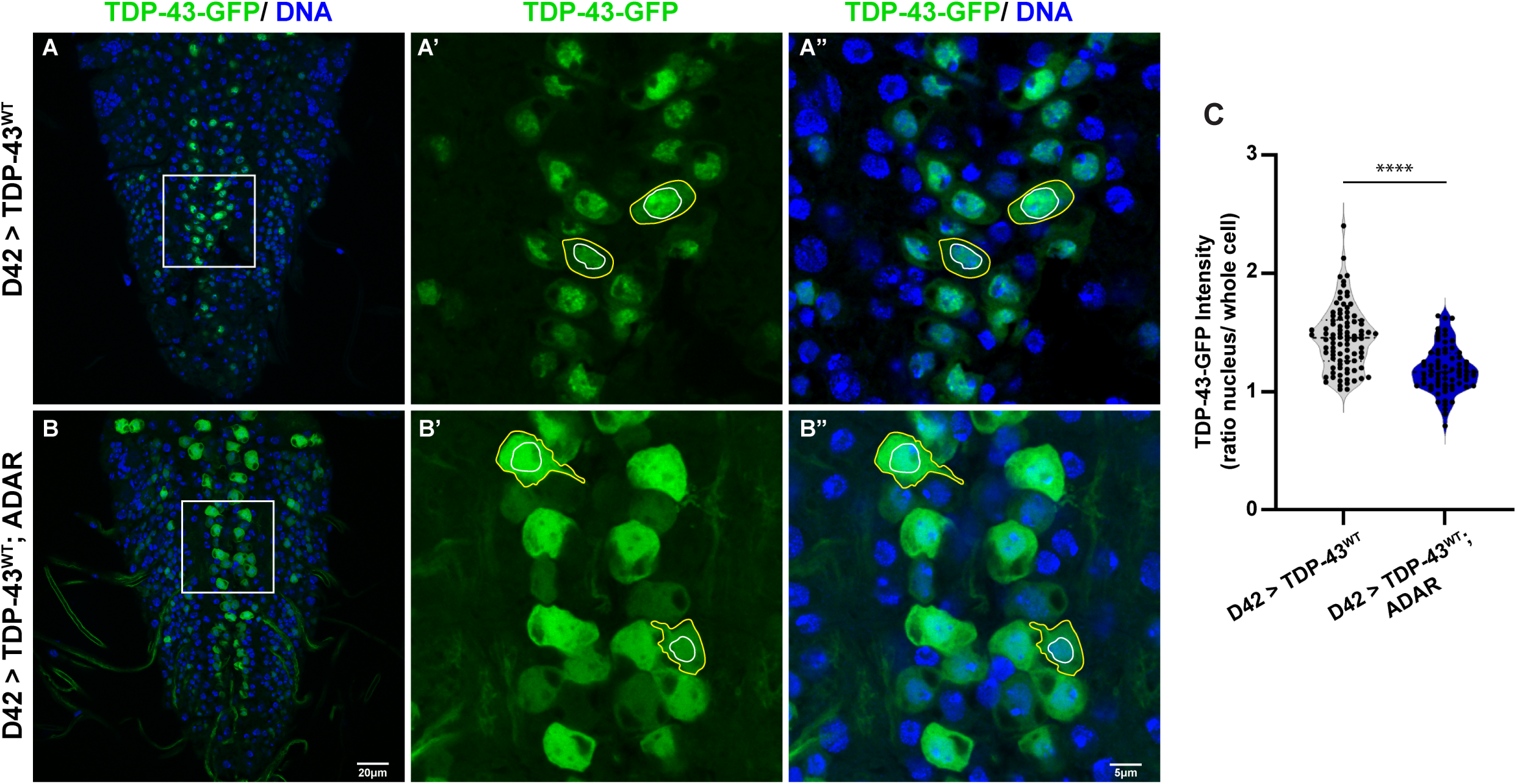
*Drosophila* motor neuron TDP-43 preferentially binds A-to-I edited RNA transcripts and is mislocalized upon *dADAR* co-expression **a)** Overexpression of ADAR increases TDP-43 cytoplasmic mislocalization in *Drosophila* motor neurons within the 3^rd^ instar larval VNC. Laval motor neurons stained for TDP-43-YFP (green) and nuclei (blue) in the VNC of A) D42 > hTDP-43^WT^ (N VNCs = 10; N cells = 90). A’, A” highlight the neurons in a zoomed-in area of the VNC. Cytoplasm area (whole cells), outlined with a yellow line; nucleus area, outlined with a white line. **b)** D42 > hTDP-43^WT^; ADAR (N=6; N cells = 70). White line square, zoomed-in area. B’ B” highlight the neurons in a zoomed-in area of the VNC. Cytoplasm area (whole cells), outlined with a yellow line; nucleus area, outlined with a white line. **c)** Quantification of TDP-43-GFP nuclear to cytoplasmic ratio. ****p=0.0001, Welch’s t test. Scale bars: B: 20µm; B”:5µm.

Together, these findings demonstrate that ADAR2 induces TDP-43 mislocalization through a previously unrecognized mechanism that requires both RNA editing activity and RNA binding to TDP-43. This effect is conserved across *in vitro* and *in vivo* systems, highlighting RNA editing as a novel regulator of TDP-43 localization.

### Increased ADAR2 activity drives TDP-43 mislocalization through nuclear export

To test whether increased A-to-I RNA editing directly alters TDP-43 nuclear export, rather than altering cytoplasmic TDP-43 aggregation, we exploited an interspecies heterokaryon assay, which has been used previously in the context of nuclear TDP-43 export (34,35,56). In this assay, SH-SY5Y cells in which endogenous TDP-43 is labeled at the C-terminus with eGFP (36) were used as a donor cell and mouse embryonic fibroblasts (MEFs) as the recipient cell. This cellular model also enabled us to test ADAR2-mediated TDP-43 mislocalization in the context of endogenous TDP-43 expression, avoiding the potential contributing aspects of TDP-43 overexpression.

SH-SY5Y donor nuclei were identified by GFP signal and positive immunostaining for the human-specific protein HNRNPC, whereas MEF nuclei were distinguished by punctate Hoechst staining and absence of HNRNPC signal (Figure 3A,B). Following protein synthesis inhibition and cell fusion, heterokaryons were defined as single fused cells exhibiting continuous wheat germ agglutinin (WGA) membrane staining and containing one donor SH-SY5Y nucleus and one recipient MEF nucleus (Figure 3C). Donor nuclei were identified by HNRNPC positivity or diffuse Hoechst staining, while recipient nuclei were defined by lack of HNRNPC signal and punctate Hoechst staining. Cells containing more than two nuclei were excluded from analysis to avoid variability associated with multinucleated heterokaryons.

**Figure 3.**
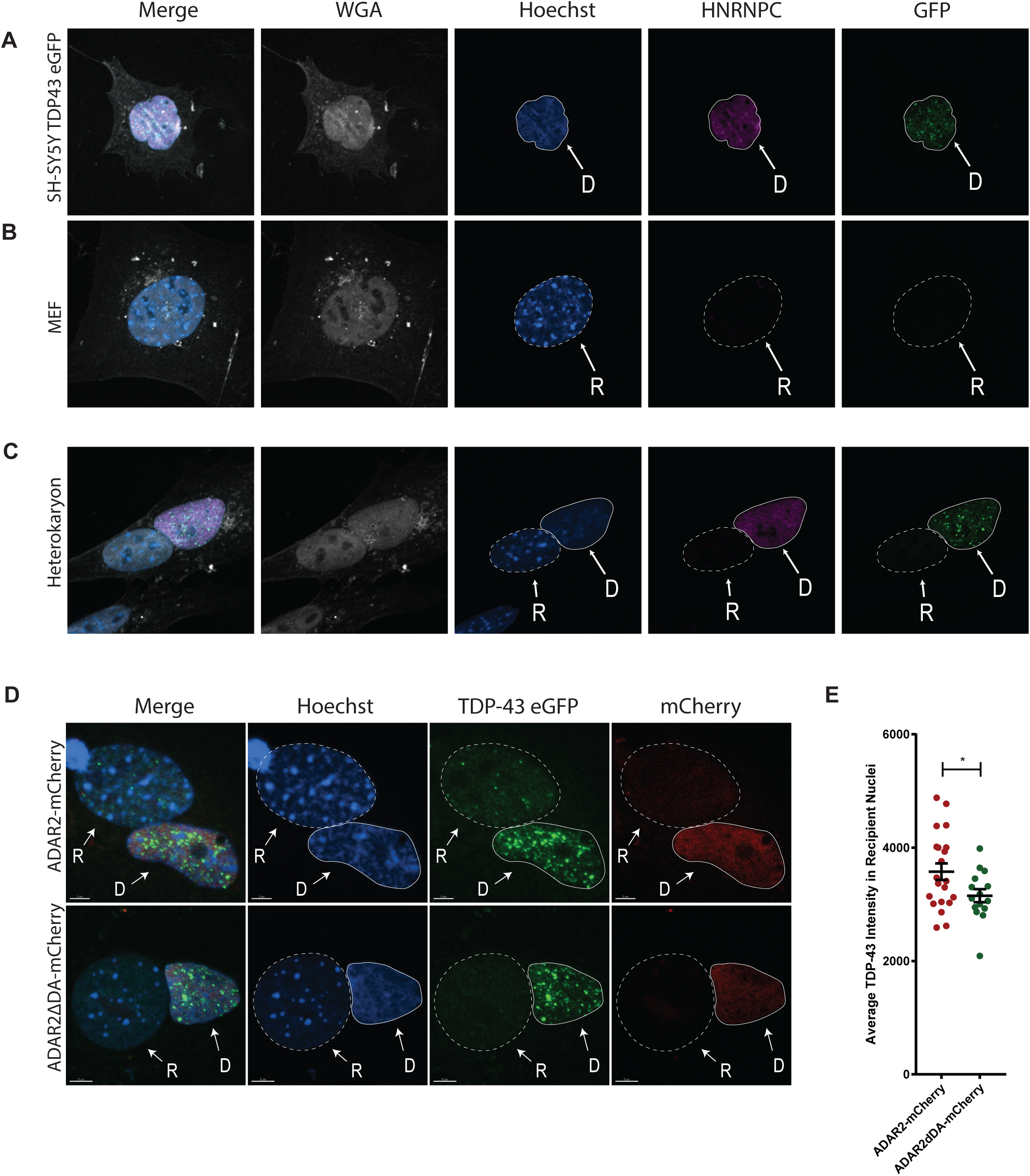
TDP-43 nuclear export is driven by ADAR2-mediated editing activity in an interspecies heterokaryon assay **a)** Representative immunofluorescence images from SH-SY5Y cells with endogenous GFP-tagged TDP-43 (TDP43-eGFP) that are used as Donor (D) cells in the heterokaryon assay. SH-SY5Y TDP43-eGFP cells exhibit diffuse Hoechst labeling, are positive for HNRNPC, and exhibit endogenous GFP signal. **b)** Representative immunofluorescence images from mouse embryonic fibroblasts (MEFs) that are used as Recipient (R) cells in the heterokaryon assay. MEFs exhibit punctate Hoechst labeling, and stain negative for HNRNPC and any endogenous GFP signal. **c)** Interspecies heterokaryons are defined by: a continuous cytoplasm, indicated by WGA staining; one donor (D) nuclei with diffuse Hoechst, which is positive for human HNRNPC; and one recipient (R) nuclei with punctuated Hoechst, which is negative for human HNRNPC. **d)** Representative images of heterokaryons formed with MEFs and SH-SY5Y TDP43-eGFP cells transduced with ADAR2-mCherry or ADAR2ΔDA-mCherry lentiviral constructs following 3h of shuttling. **e)** Heterokaryons with ADAR2-mCherry overexpression have more endogenous TDP-43-eGFP detected in the recipient MEF nuclei than heterokaryons with ADAR2ΔDA-mCherry overexpression. (N = 15 heterokaryons analyzed, *p< 0.05, t-test, scale bar = 2µm)

To validate this assay for quantifying endogenous TDP-43 nuclear export, we first examined a series of post-fusion shuttling intervals (Supplementary Figure 2). Quantification of the TDP-43 shuttling index demonstrated progressive nuclear export after 3, 6, 9, and 12 h (Supplementary Figure 3A; p < 0.0001), whereas no change was observed at earlier time points (0.5 - 1 h), confirming that this model system enables quantitative measurement of endogenous TDP-43 nuclear export.

We next used this approach to test whether enhanced ADAR2 activity influences TDP-43 export. SH-SY5Y donor cells were transduced with lentiviral constructs expressing either wild-type ADAR2-mCherry or catalytically inactive ADAR2ΔDA-mCherry prior to fusion with recipient MEFs. Heterokaryons formed from SH-SY5Y cells transduced with wild-type ADAR2 exhibited increased TDP-43-eGFP signal in recipient MEF nuclei, whereas minimal signal was observed in heterokaryons formed with ADAR2ΔDA transduction (Figure 2D,E; p < 0.05).

Together, these results indicate that ADAR2 catalytic activity promotes nuclear export of TDP-43, contributing to its cytoplasmic mislocalization.

### UI-repeat RNA oligomers are sufficient to facilitate TDP-43 nuclear export

Having established that overexpression of catalytically active ADAR2 is sufficient to trigger TDP-43 nuclear export, we wondered whether A-to-I edited RNA oligomers alone could influence nuclear TDP-43 trafficking. Using an established nuclear export assay (40), permeabilized HeLa cells were incubated for 30 minutes with varying RNA oligomer motifs at multiple concentrations, followed by immunostaining for TDP-43 (Figure 4A). The oligomers consisted of twelve nucleotides each and resembled either: adenosines only ((A)_12_); a mix of un-edited adenosines and uracils ((AU)_6_); an “A-to-I” edited version of AU12 ((IU)_6_); or TDP-43’s ideal binding motif of GU-rich repeats ((GU)_6_). Incubation with (AU)_6_ or (A)_12_ oligomers demonstrated no change in TDP-43 nuclear intensity relative to untreated HeLa cells across all concentrations (Figure 4B,C; p>0.05). However, similar to (GU)_6_ oligomers, representing TDP-43’s binding motif, (IU)_6_ resulted in a concentration-dependent loss of nuclear TDP-43 signal (Figure 4B,C; p<0.001, p<0.01, respectively). While (GU)_6_ oligomers promoted a slightly larger quantifiable drop in TDP-43 nuclear intensity, the drastic loss of nuclear TDP-43 signal with the (IU)_6_ oligomers highlights the ability of an A-to-I edited RNA transcript to alter the localization of TDP-43 within the cell in support of our hypothesis.

**Figure 4.**
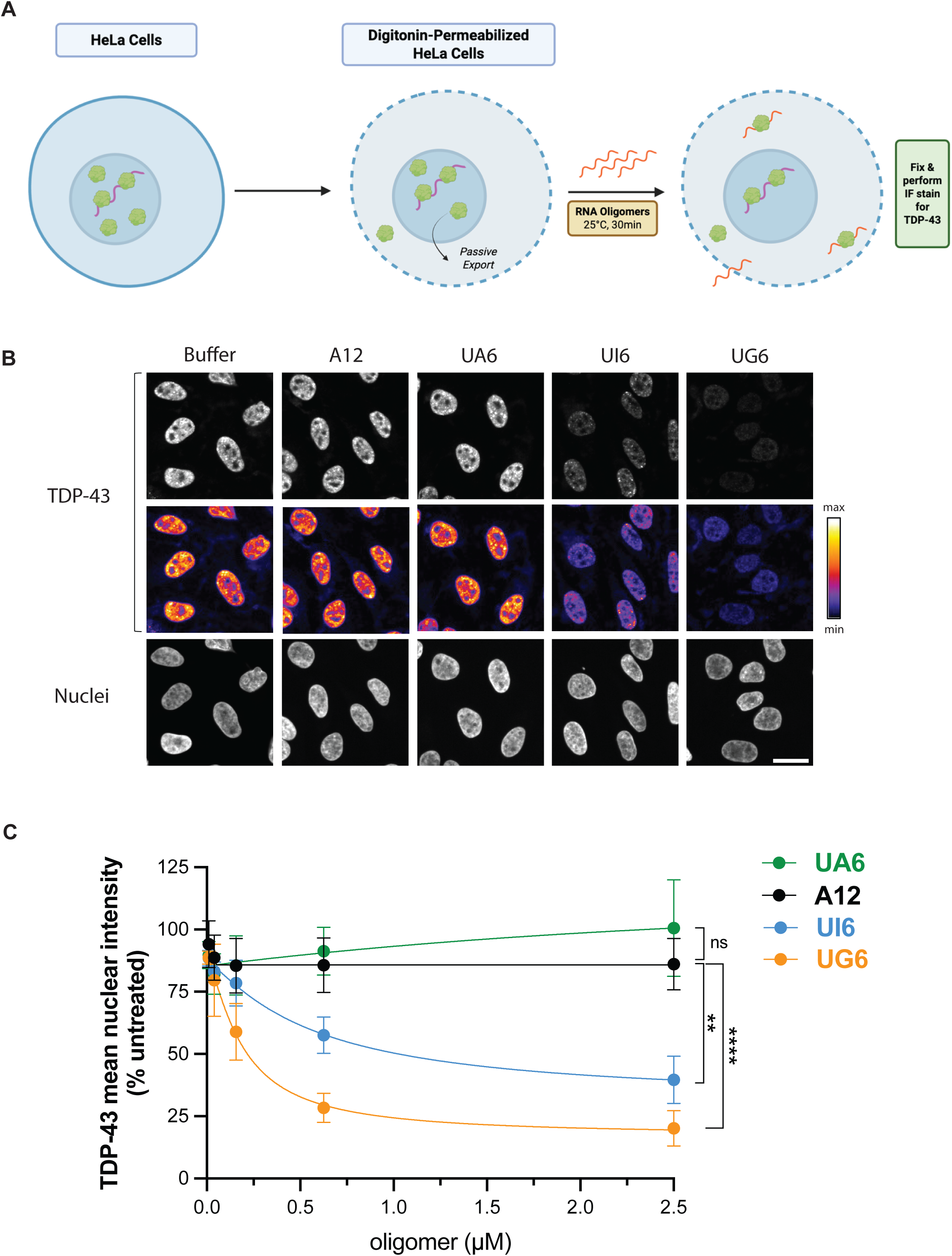
“Edited” RNA oligomers promote TDP-43 nuclear export in a HeLa cell assay **a)** Schematic detailing the nuclear export assay in permeabilized HeLa cells, created with institutional license at Biorender.com **b)** Immunofluorescence images of nuclear TDP-43 intensity counterstained with DAPI; scale bar = 25µm. **c)** Quantification of nuclear intensity of TDP-43 relative to untreated cells for each RNA oligomer at multiple concentrations (0.5-2.5µM); N=4 repeated experiments, >3,000 cells per condition per experiment; **p<0.01, ****p<0.001, one-way ANOVA.

### Inosine-containing RNAs bind to TDP-43 and are enriched in TDP-43-bound transcripts *in vivo*

Based on our findings that ADAR2 activity influences TDP-43 localization, we hypothesized that A-to-I RNA editing may alter TDP-43 RNA interactions by generating or modifying binding motifs. To test whether inosine-containing RNAs can directly interact with TDP-43, we performed electrophoretic mobility shift assays (EMSAs) using recombinant TDP-43 and synthetic RNA oligonucleotides. We compared binding to canonical UG repeats ((UG)₁₂), as well as unedited ((UA)₁₂) and inosine-containing ((UI)₁₂) sequences. As expected, TDP-43 exhibited high affinity for UG repeats (Kd ∼ 0.1 µM)∫. In contrast, binding to (UA)₁₂ was weak (Kd ∼ 100 µM), whereas (UI)₁₂ displayed a ∼50-fold increase in affinity compared to (UA)₁₂ (Kd ∼ 2 µM) (Figure 5A–F). Interestingly, mutant TDP-43 harboring the R151A point mutation that disrupts intra-molecular salt bridges in TDP-43 and subsequently impairs RNA binding to TDP-43 (38) also exhibited a reduction in binding to (UI)₁₂ compared to (UG)₁₂ (Supplementary Figure 4). These EMSA results indicate that inosine incorporation can indeed enhance TDP-43 binding relative to unedited RNA, although with lower affinity than canonical UG-rich motifs.

**Figure 5.**
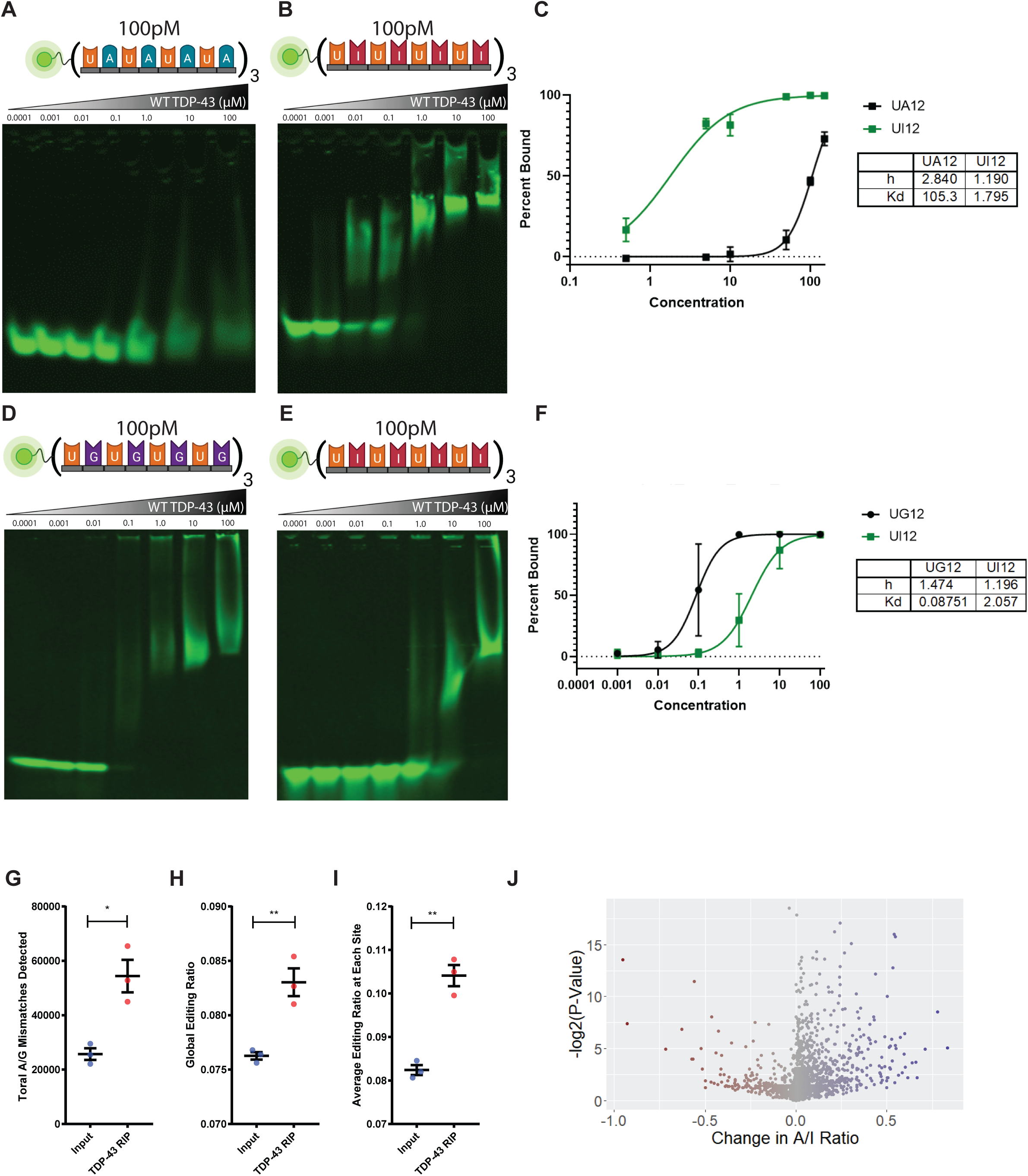
TDP-43 exhibits cooperative binding to RNAs containing GU- and IU-rich repetitive regions **a)** Increasing concentrations of recombinant WT TDP-43 (0.5 µ M – 150 µM) were mixed with labeled (UA)_12_ (100pM). **b)** Increasing concentrations of recombinant WT TDP-43 (0.001 µ M – 100 µM) were mixed with labeled (UI)_12_ (100pM). **c)** Calculations and plotting of Kd and h for (UA)_12_ and (UI)_12_; n=3 different replicates. **d)** Increasing concentrations of recombinant WT TDP-43 (0.5 µ M – 150 µM) were mixed with labeled (UG)_12_ (100pM). **e)** Increasing concentrations of recombinant WT TDP-43 (0.001 µ M – 100 µM) were mixed with labeled (UI)_12_ (100pM). **f)** Calculations and plotting of Kd and h for (UG)_12_ and (UI)_12_; n=3 different replicates. **g)** Analysis of TDP-43 RIP-seq data reveals the total number of A/G mismatches is increased in RNA bound to TDP-43 in *Drosophila* motor neurons. N=3 replicates, 100 flies per replicate, *p<0.05, Student’s t-test. **h)** The Global RNA Editing Ratio of TDP-43-bound mRNA in *Drosophila* motor neurons is increased. N=3 replicates, 100 flies per replicate, **p<0.005, Student’s t-test. **i)** The average RNA editing ratio across all RNA editing sites is increased in RNA bound to TDP-43. N=3 replicates, 100 flies per replicate, **p<0.005, Student’s t-test **j)** Volcano plot demonstrating increases in most individual editing sites within TDP-43 RIP fraction relative to Input.

To determine whether this interaction is reflected in vivo, we analyzed published RIP-seq datasets from *Drosophila* larvae expressing human TDP-43-YFP in motor neurons (27) for known *Drosophila* RNA editing sites cataloged in the REDI portal (28). This analysis demonstrated that TDP-43–associated RNAs exhibited increased mismatch frequencies relative to input RNA, consistent with elevated A-to-I editing (Figure 5G; p < 0.05). Quantification of both global and site-specific editing levels revealed higher editing ratios in the TDP-43 immunoprecipitated fraction compared to input RNA (Figure 5H,I,J; p < 0.005).

Together, these findings demonstrate that inosine-containing RNAs can preferentially associate with TDP-43, supporting a model in which A-to-I RNA editing may alter the RNA binding landscape of TDP-43. While these data suggest that edited RNAs may contribute to TDP-43 redistribution, they do not exclude the possibility that editing may also disrupt existing binding interactions, indicating that ADAR2 activity and RNA editing could modulate TDP-43 function through multiple mechanisms.

### RNAseq and eCLIP-seq identify editing-associated changes in TDP-43 RNA binding

Collectively, our results thus far suggest that increased ADAR2 activity alters TDP-43 localization across multiple cell types and model systems. However, whether enhanced A-to-I RNA editing also modifies the broader RNA interaction landscape of TDP-43 remained unclear. To address this, we generated SH-SY5Y cell lines stably expressing doxycycline-inducible ADAR2 variants with differing editing activities: wild-type ADAR2 (ADAR2^WT^), a hypoactive mutant (ADAR2^E396A^) and a hyperactive mutant (ADAR2^E488Q^). Following doxycycline induction, we performed bulk RNA sequencing (RNAseq) together with TDP-43 enhanced cross-linking and immunoprecipitation followed by RNA sequencing (eCLIP-seq) to assess editing-depenent changes in gene epression and TDP-43 RNA binding substrates (Figure 6A) (41).

**Figure 6.**
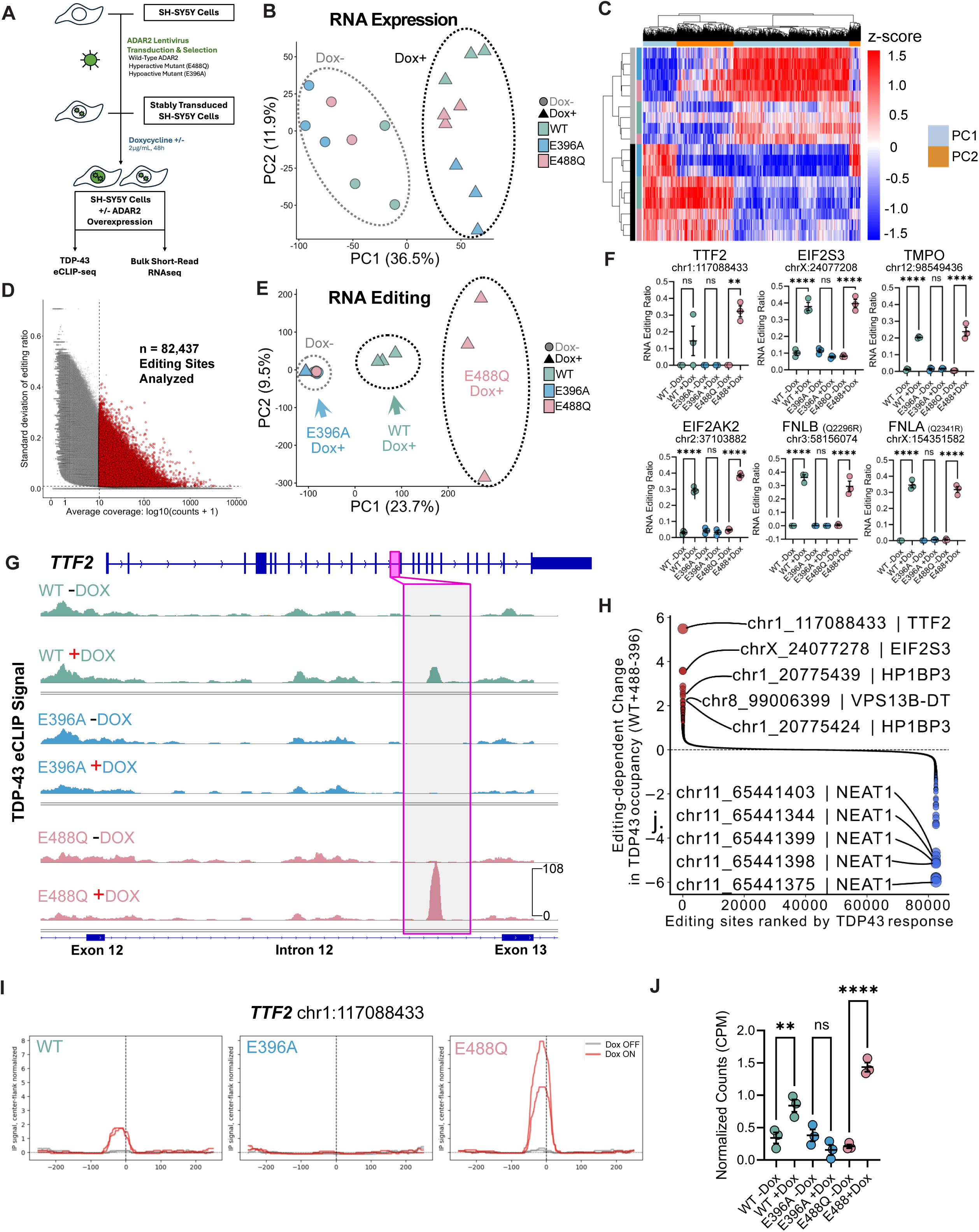
ADAR2 editing activity-dependent shifts in TDP-43’s RNA binding profile are detected via eCLIP-seq **a)** Experimental schematic of ADAR2 lentivirus transduction, Doxycycline induction, and eCLIP-seq and RNAseq workflows. **b)** PCA plot between Dox+ and Dox-conditions for each ADAR2 lentivirus group. **c)** Heatmap of differentially expressed RNA transcripts grouped by editing activity and Dox+/- condition. **d)** Scatterplot of 82,437 A-to-I RNA editing sites analyzed based on coverage and editing ratio. **e)** PCA plot of RNA editing sites grouped by editing activity and Dox+/- condition. **f)** Comparisons of editing ratios between TTF2, EIF2S, TMPO, EIF2AK2, FNLA, and FNLB between Dox+/- conditions. One-way ANOVA, **p<0.01, ****p<0.001, ns = not significant. **g)** Highlight of differential TDP-43 eCLIP-seq peaks in TTF2 mediated by active ADAR2 editing. **h)** Plot of editing-dependent differential increases and decreases in TDP-43 binding sites with active ADAR2 overexpression. **i)** Plots of TDP-43 binding sites in TTF2 exhibiting differential editing in ADAR2 WT and E488Q overexpression, but not E396A overexpression. **j)** ANOVA of normalized counts per million for TTF2 cryptic exon inclusion between Dox+/- conditions for ADAR2 lentiviruses. One-way ANOVA, **p<0.01, ****p<0.001, ns = not significant.

Principal component analysis (PCA) of bulk RNAseq data revealed that ADAR2 induction represented the dominant source of transcriptional variance across samples (PC1, 36%; Figure 6B). Differential expression analysis further identified 1,565 transcripts altered in an editing-dependent manner following ADAR2 overexpression (Figure 6C, Supplemental Dataset 1). These findings indicate that enhanced ADAR2 activity broadly influences the transcriptional landscape of SH-SY5Y cells.

We next wondered whether induction of active ADAR2 variants altered cellular RNA editing profiles. Following filtering based on editing ratios and coverage (Figure 6D), PCA of identified editing sites demonstrated clear separation of ADAR2^WT^ and ADAR2^E488Q^ doxycycline-induced samples from both ADAR2^E396A^ Dox+ and non-induced conditions (PC1, 23.7%; Figure 6E). This findings confirm that induction of catalytically active ADAR2 variants produces distinct RNA editing signatures, while the inactive point mutation E396A does not substantially alter the endogenous editing landscape.

To identify editing-dependent TDP-43 RNA interactions, we integrated RNA editing analyses with our TDP-43 eCLIP-seq dataset. This approach identified 7,803 RNA editing sites across 2,700 genes that were associated with altered TDP-43 occupancy under conditions of increased editing activity (Supplemental Dataset 2). Representative examples included intronic editing sites in *TTF2, EIF2S3, TMPO, and EIF2AK2* as well as exonic editing sites predicted to generate glutamine-to-arginine (Q-R) substitutions in *FNLB* and *FNLA* (Figure 6F).

We next quantified RNA editing-dependent changes in TDP-43 binding by comparing TDP-43 eCLIP-seq across RNA editing conditions (ADAR2 WT and ADAR2^E488Q^ Dox+). While relatively few editing sites demonstrated increased TDP-43 occupancy following enhanced editing activity (Figure 6H), the most prominent example corresponded to an edited *Alu* repetitive element located within intron 12 of the transcription termination factor 2 (*TTF2*) transcript (Figure 6G, I). Increased TDP-43 binding at this locus was observed specifically in ADAR2^WT^ and ADAR2^E488Q^ doxycycline-induced cells, coinciding with the appearance of a *TTF2* cryptic exon that was absent in ADAR2^E396A^-expressing cells (Figure 6J). These findings suggest that increased RNA editing can remodel local TDP-43 binding patterns and may contribute to altered transcript processing at specific loci. In contrast, a larger subset of editing sites exhibited reduced TDP-43 occupancy under conditions of elevated editing activity. Notably, several of the most prominent negatively-associated sites mapped to the long non-coding RNA *NEAT1* (Figure 6H), suggesting that increased RNA editing may disrupt TDP-43 interactions with transcripts involved in nuclear RNA regulatory pathways.

Together, these findings suggest that increased ADAR2 activity not only alters global RNA editing patterns and transcriptional profiles, but also reshapes the RNA-binding landscape of TDP-43 in an editing-dependent manner. These data support a model in which A-to-I RNA editing modifies TDP-43 interactions with specific RNA substrates, potentially contributing to altered TDP-43 localization and downstream RNA dysregulation.

## DISCUSSION

TDP-43 nuclear depletion and cytoplasmic accumulation are defining pathological features of ALS, FTD, and multiple related neurodegenerative disorders, yet the molecular events that initiate these changes remain incompletely understood (5,7,57). Post-transcriptional RNA modifications, such as methylation of adenosine N6 (m6A modifications), can alter TDP-43’s binding preference for methylated RNA, possibly due to altering the accessibility of TDP-43 motifs for proper binding (52). Here, we identify A-to-I RNA editing as a previously unrecognized regulator of TDP-43 localization and RNA interactions. Across multiple orthogonal model systems, we demonstrate that increased ADAR2-mediated RNA editing promotes redistribution of TDP-43 from the nucleus to the cytoplasm in a manner dependent on both ADAR2 catalytic activity and TDP-43 RNA-binding capacity. Furthermore, we show that inosine-containing RNAs are sufficient to enhance TDP-43 nuclear export and can directly alter TDP-43 RNA-binding interactions. Together, these findings support a model in which dysregulated RNA editing alters TDP-43–RNA interactions and contributes to abnormal TDP-43 nucleocytoplasmic trafficking.

Previous studies assessing ADAR2 dysfunction in neurodegenerative diseases have primarly focused on the loss of ADAR2 function, partiularly in the context of altered GluA2 editing and excitotoxicity in ALS and in ALS/AD (58–62). Furthermore, our previous studies exploring ADAR2 in C9-ALS/FTD noted mislocalization and cytoplasmic accumulation of ADAR2 in both iPSC motor neurons and postmortem tissue, where we observed both a decrease and an increase in the A-I RNA editing profile of these tissue samples (22). In support of these studies, our findings suggest that increased or dysregulated RNA editing activity may indeed influence disease-relevant cellular processes by altering RNA-protein interactions. Importantly, our data do not argue that elevated ADAR2 activity alone is sufficient to fully recapitulate TDP-43 proteinopathy in vivo. Rather, they support the idea that altered RNA editing environments can modulate TDP-43 localization and RNA binding, potentially contributing to early pathogenic events in susceptible neuronal populations.

Our *Drosophila* studies further support the relationship between RNA editing and TDP-43 localization *in vivo*. Overexpression of dADAR promoted cytoplasmic redistribution of human TDP-43 in motor neurons, while analysis of TDP-43 RIP-seq datasets demonstrated enrichment of edited transcripts within TDP-43-associated RNAs. These findings are particularly intriguing given the established importance of dADAR in neuronal excitability, synaptic transmission, and neurodegeneration in flies (63,64). Furthermore, *dADAR* is crucial for maintenance of synaptic transmission as knockout flies have drastically reduced neurotransmitter release and alterations in activity at their neuromuscular junctions, further linking RNA editing activity to synaptic dysfunction and possible neuromuscular issues (65,66). To date and to our knowledge, the only study examining *dADAR* overexpression in flies demonstrated lethality at larval and pupal stages of development, and showed that overactive RNA editing reduced motor neuron activity while knockout of *dADAR* led to motor neuron hyperexcitability (67). Together, these observations raise the possibility that altered RNA editing may broadly influence RNA-binding protein behavior within neurons.

Recent studies have indicated that the localization of TDP-43 is dependent on binding to RNAs containing the GU-repeat motif, which is demonstrated in our nuclear export assay (40). In previous studies using computational modeling of healthy and mutant TDP-43 interactions, it was discovered that intermolecular van der Waals interactions between polar and nonpolar regions of RNA nucleotides and TDP-43 RRMs are important for proper binding and splicing properties of TDP-43; it is possible that inosine-containing transcripts do not afford as strong of an interaction with TDP-43 guanosine nucleotides, but this remains to be investigated (68). One limitation of our study is that we did not use “partially edited” oligomers to more realistically simulate an isolated A-to-I editing event.

Finally, our RNAseq and eCLIP-seq experiments reveal that ADAR2 overexpression can alter gene expression as well as create and disrupt TDP-43 binding sites in an editing activity-dependent manner. To our knowledge, these findings are the first to explore the transcriptional landscape of A-to-I edits in the context of TDP-43 binding. We identified editing-dependent changes in both gene expression and TDP-43 binding sites, suggesting that altered RNA editing may redirect TDP-43 interactions across the transcriptome. The act of RNA modifications altering the binding of TDP-43 and other RBPs has been previously described, providing support for our hypothesis (52,69–72). Particularly interesting was the identification of editing-associated changes in TDP-43 binding within transcripts linked to neurodegeneration and RNA regulation, including *FLNB, FNLA*, *TTF2*, and *NEAT1*. While the functional significance of these individual interactions remains to be determined, these findings suggest that altered RNA editing could influence multiple downstream pathways through redistribution of TDP-43 RNA binding. A recent study has implicated *FLNB* as a misspliced RNA in *C9ORF72* patient-derived iPSC neurons (73). The transcript showing the strongest editing-dependent increase in TDP-43 is *TTF2*, which is an ATP-dependent repressor of transcription during mitosis, which may imply there is regulation of cell cycle dynamics in an environment of increased RNA editing (74). Fascinatingly, *NEAT1* transcripts exhibit a reduction in TDP-43 binding when RNA editing is artificially increased within our SH-SY5Y cells. *NEAT1* is known to increase in response to rising TDP-43 levels to reduce the neurotoxic effects of TDP-43 proteinopathy in yeast and *Drosophila*, while the loss of *NEAT1* in response to TDP-43 loss is responsible for increased degeneration of motor neurons(75,76). The consequences of altered TDP-43 binding to these transcripts remains an area of active interest and investigation.

Several limitations of the current study should be acknowledged. First, many experiments relied on ADAR2 overexpression paradigms, which may not fully recapitulate endogenous editing dynamics observed in disease. Future studies will be required to determine whether endogenous alterations in ADAR activity or transcript-specific editing changes are sufficient to produce similar TDP-43 redistribution in disease-relevant neuronal systems. Second, while our data support a relationship between edited RNAs and TDP-43 nuclear export, the precise molecular mechanisms linking RNA editing to nucleocytoplasmic trafficking remain unresolved. Third, our sequencing analyses identify editing-dependent changes in TDP-43 binding, but additional functional studies will be necessary to determine which specific edited transcripts contribute directly to TDP-43 mislocalization or downstream toxicity. Importantly, ADAR proteins also exert multiple editing-independent functions, including miRNA regulation, RNA splicing, retrotransposon suppression, and expression of proteins involved in the innate immune response (77–80). Although our catalytically inactive ADAR2 controls support an editing-dependent mechanism in the present study, we cannot exclude the possibility that editing-independent ADAR functions additionally contribute to altered TDP-43 biology under pathological conditions.

## CONCLUSION

Overall, our findings identify A-to-I RNA editing as a novel regulator of TDP-43 localization and RNA interactions. More broadly, this work highlights how post-transcriptional RNA modifications may influence RNA-binding protein behavior and nucleocytoplasmic trafficking in neurodegenerative disease. Given the widespread involvement of TDP-43 pathology across ALS, FTD, Alzheimer’s disease, and related disorders, understanding how RNA editing environments shape TDP-43 function may provide new insight into early pathogenic mechanisms and reveal new therapeutic opportunities targeting RNA regulatory pathways.

## LIST OF ABBREVIATIONS

A-to-I: Adenosine to inosine
ADAR: Adenosine deaminase acting on RNA
ADAR2: Adenosine deaminase acting on RNA 2
ALS: Amyotrophic lateral sclerosis
C9: *C9ORF72,* Chromosome 9 open reading frame 72
C9 ALS/FTD: *C9ORF72*-associated ALS and FTD
dADAR: *Drosophila* ADAR
eCLIP-seq: Enhanced cross-linking and immunoprecipitation followed by RNA sequencing
eGFP: Endogenous green fluorescent protein
EMSA: Electrophoretic mobility shift assays
FTD: Frontotemporal dementia
LATE: Limbic-predominant age-related TDP-43 encephalopathy
MEF: Mouse embryonic fibroblasts
N:C: Nucleocytoplasmic
RIP: RNA immunoprecipitation
RIP-seq: RNA immunoprecipitation followed by RNA sequencing
RNAseq: RNA sequencing
TDP-43: Transactive response DNA binding protein of 43 kilodaltons
GFP: Green fluorescent protein
UMI: Unique molecular identifier
WGA: Wheat germ agglutinin
YFP: Yellow fluorescent protein

## DECLARATIONS

- Ethics approval and consent to participate

- Not applicable.
- Consent for publication

- Not applicable.
- Availability of data and materials

- The RNAseq and eCLIP-seq datasets generated during the current study will be available in the GEO repository, with a link to be uploaded upon acceptance.
- The RIP-seq datasets analyzed during the current study are available in the GEO repository, https://www.ncbi.nlm.nih.gov/geo/query/acc.cgi?acc=GSE156222 (27)
- Competing interests

- The authors declare that they have no competing interests.
- Funding

- NIH R21NS130492 to RS
- NIH/NINDS R01NS097542 and 2R37NS113943-06, NIH/NIA P30 AG072931 to SJB
- NIH R01NS091299 to DCZ
- TDP-43 eCLIP-seq was financially supported by a RBP-eCLIP Matching Fund Award from Eclipse Biosciences awarded to DLJ and RS.
- Authors’ Contributions

- HEK293 experiments and data analyses were performed by SM and DLJ.
- Heterokaryon assays and data analyses were performed by SM, LMG, and IL.
- Drosophila experiments and data analyses were performed by EL, SM, DM, and DCZ.
- HeLa cell experiments and data analyses were performed by PK and LH.
- EMSA experiments were performed by MM and SJB.
- Drosophila RIPseq analyses, RNAseq analyses and A-to-I editing analyses were performed by SM, EA and KVK.
- SH-SY5Y ADAR2 experiments were performed by SM and DLJ.
- RBP-eCLIP-seq experiments and sample preparation were performed by DLJ.
- SH-SY5Y RNAseq and eCLIP-seq combinatorial analyses were performed by SM and EA.
- Figure generation was performed by SM and DLJ.
- SM, DLJ and RS were the primary writers of the manuscript.

## Acknowledgements

- The authors thank the members of the Sattler, Barmada, Donnelly, Zarnescu, Van Keuren-Jensen labs for their assistance in experiments, sequencing, and critical reading of this manuscript.
- We thank Dr. Gene Yeo and the members of his lab for discussions and guidance on interpreting our initial eCLIP-seq results.
- We thank Dr. Don Cleveland and Dr. Fatima Gasset-Rosa for their generous contribution of eGFP-tagged SH-SY5Y cells for the heterokaryon assay (36).

## TABLES

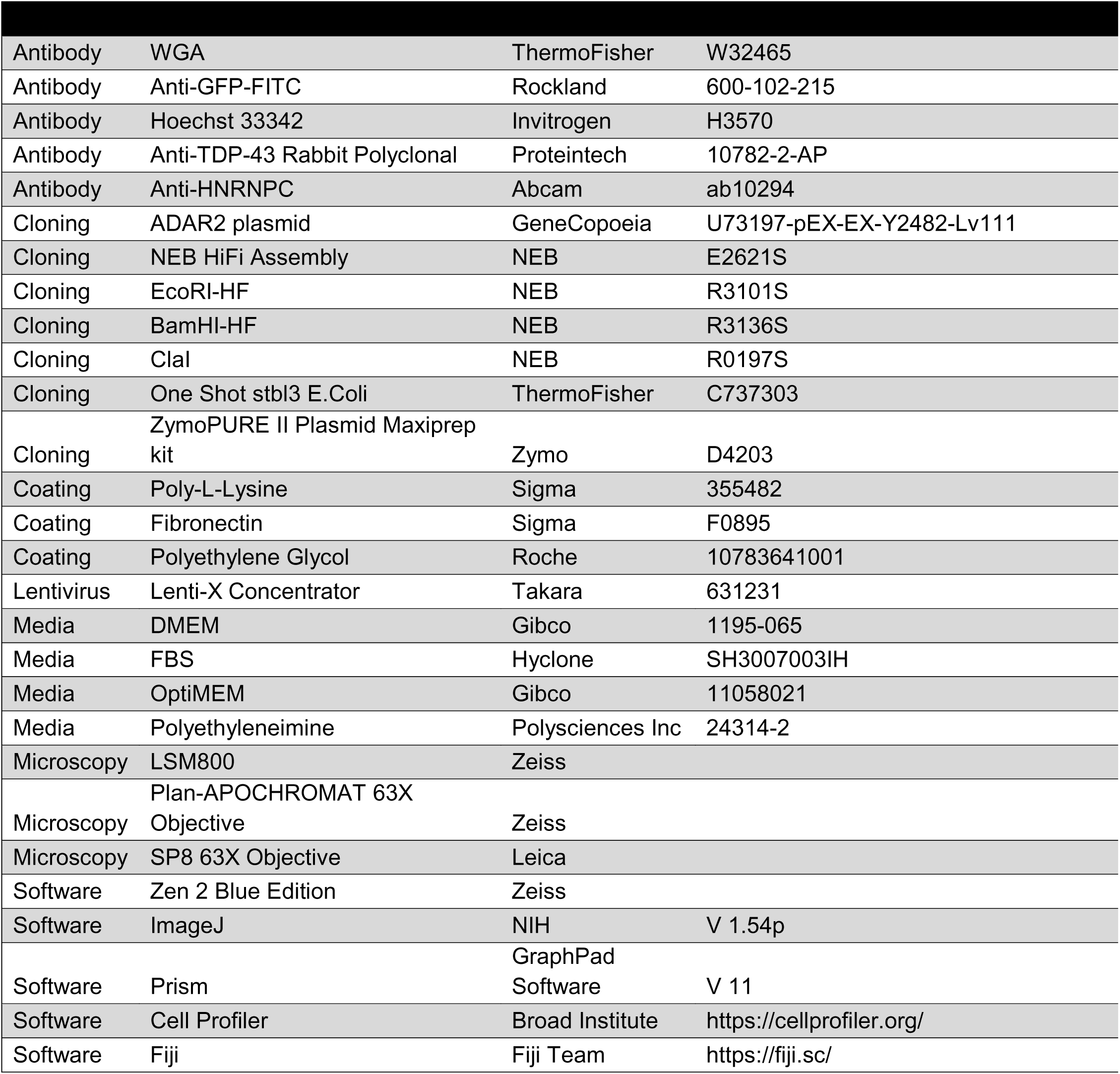

| Antibody | WGA | ThermoFisher | W32465 |
| --- | --- | --- | --- |
| Antibody | Anti-GFP-FITC | Rockland | 600-102-215 |
| Antibody | Hoechst 33342 | Invitrogen | H3570 |
| Antibody | Anti-TDP-43 Rabbit Polyclonal | Proteintech | 10782-2-AP |
| Antibody | Anti-HNRNPC | Abcam | ab10294 |
| Cloning | ADAR2 plasmid | GeneCopoeia | U73197-pEX-EX-Y2482-Lv111 |
| Cloning | NEB HiFi Assembly | NEB | E2621S |
| Cloning | EcoRI-HF | NEB | R3101S |
| Cloning | BamHI-HF | NEB | R3136S |
| Cloning | Clal | NEB | R0197S |
| Cloning | One Shot stbl3 E.Coli | ThermoFisher | C737303 |
| Cloning | ZymoPURE II Plasmid Maxiprep kit | Zymo | D4203 |
| Coating | Poly-L-Lysine | Sigma | 355482 |
| Coating | Fibronectin | Sigma | F0895 |
| Coating | Polyethylene Glycol | Roche | 10783641001 |
| Lentivirus | Lenti-X Concentrator | Takara | 631231 |
| Media | DMEM | Gibco | 1195-065 |
| Media | FBS | Hyclone | SH3007003IH |
| Media | OptiMEM | Gibco | 11058021 |
| Media | Polyethyleneimine | Polysciences Inc | 24314-2 |
| Microscopy | LSM800 | Zeiss |  |
| Microscopy | Plan-APOCHROMAT 63X Objective | Zeiss |  |
| Microscopy | SP8 63X Objective | Leica |  |
| Software | Zen 2 Blue Edition | Zeiss |  |
| Software | ImageJ | NIH | V 1.54p |
| Software | Prism | GraphPad Software | V 11 |
| Software | Cell Profiler | Broad Institute | https://cellprofiler.org/ |
| Software | Fiji | Fiji Team | https://fiji.sc/ |

