## Supplemental Figures for "ADAR2-Mediated RNA Editing Promotes TDP-43 Nuclear Export and Alters RNA Binding"

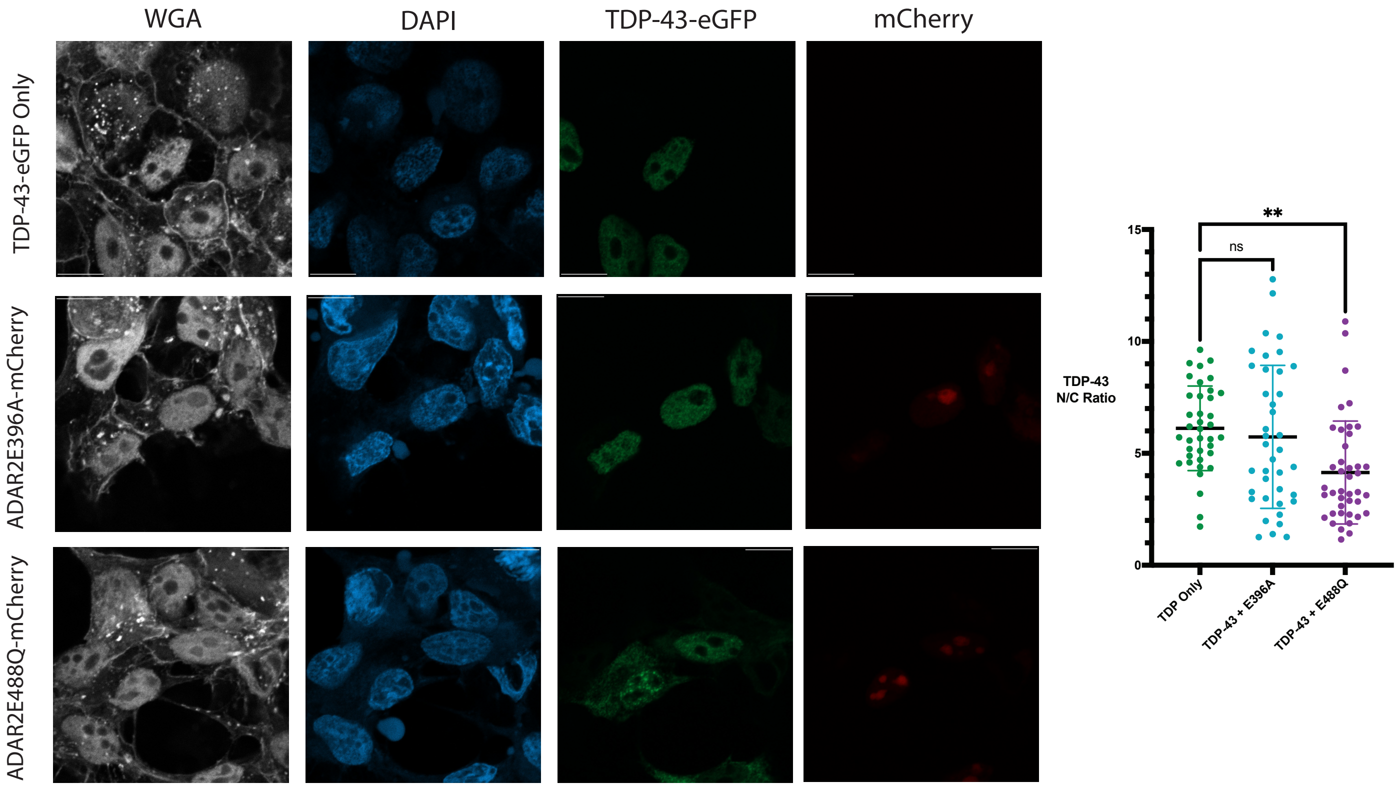


**Supplemental Figure 1: Co-transfection of TDP-43 with a hyperactive, but not a hypoactive, ADAR2 variant in HEK293T cells lowers the N/C ratio of TDP-43.**

a) Representative images of TDP-43-eGFP mislocalization with ADAR2$^{E488Q}$-mCherry overexpression but not with ADAR2$^{E396A}$-mCherry; scale bar = 10µm.

b) TDP-43-eGFP N/C Ratio is decreased in HEK293T cells overexpressing ADAR2$^{E488Q}$-mCherry; n=38, 39, and 41 cells/group for TDP Only, TDP-43 + E396A, and TDP-43 + E488Q, respectively; one-way ANOVA, **$p<0.01$, ns = not significant.

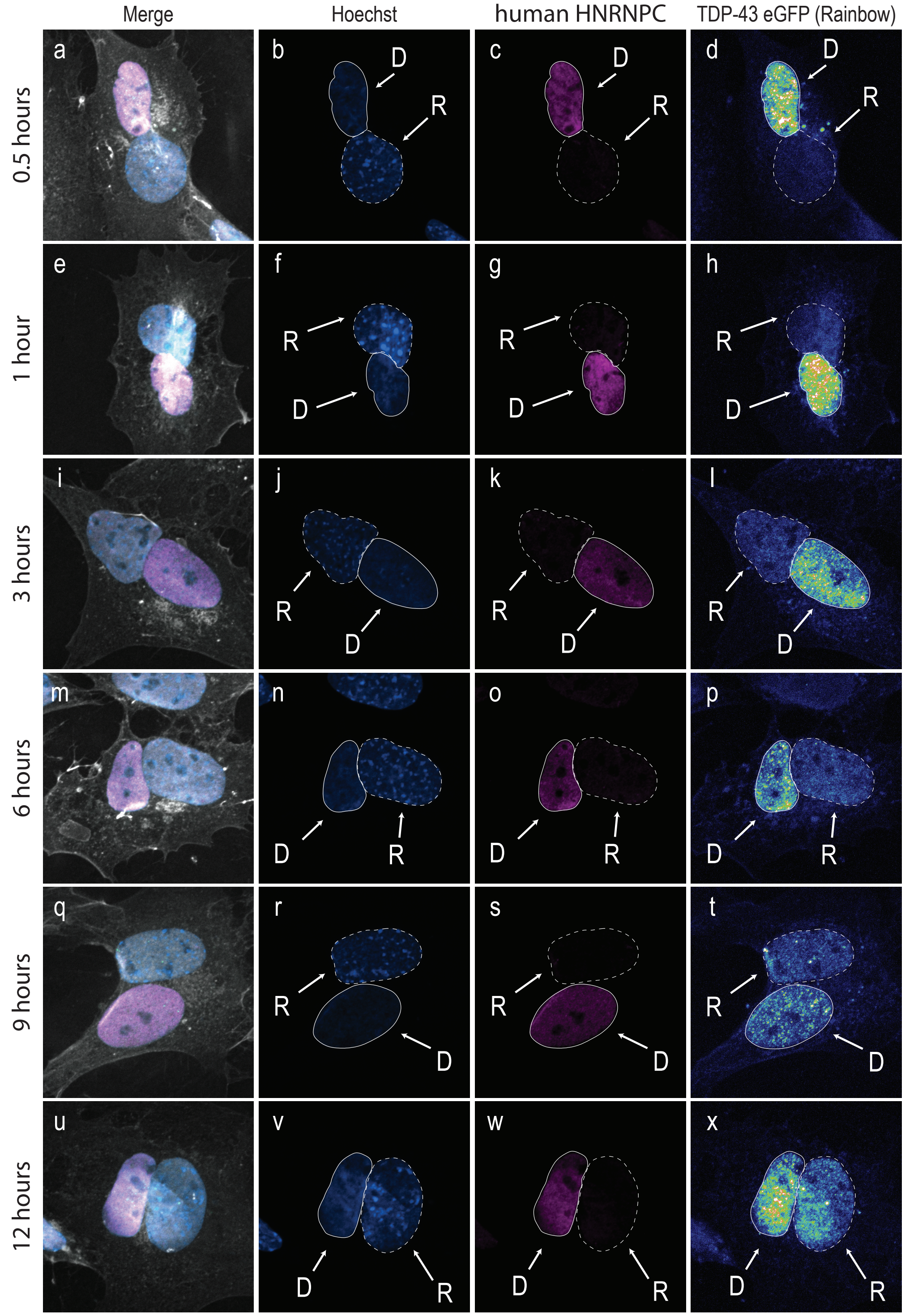


**Supplemental Figure 2: TDP-43 nuclear export can be measured across tiem with heterokaryon assays.** **a,e,i,m,q,u**: Representative images of interspecies heterokaryons that have undergone 0.5, 1, 2, 3, 6, 9, and 12 hours of shuttling time, respectively. **a-c, e-g, i-k, m-o, q-s, u-w**: Heterokaryons were selected for analysis if they had continuous WGA staining, one donor nuclei and one recipient nuclei, indicated by Hoechst and human HNRNPC labeling. **d, h, l, p, t, s**: The nuclear export of endogenous TDP-43 eGFP was assessed by the pixel intensity of regions of interest manually defined around the recipient and donor nuclei at each time point.

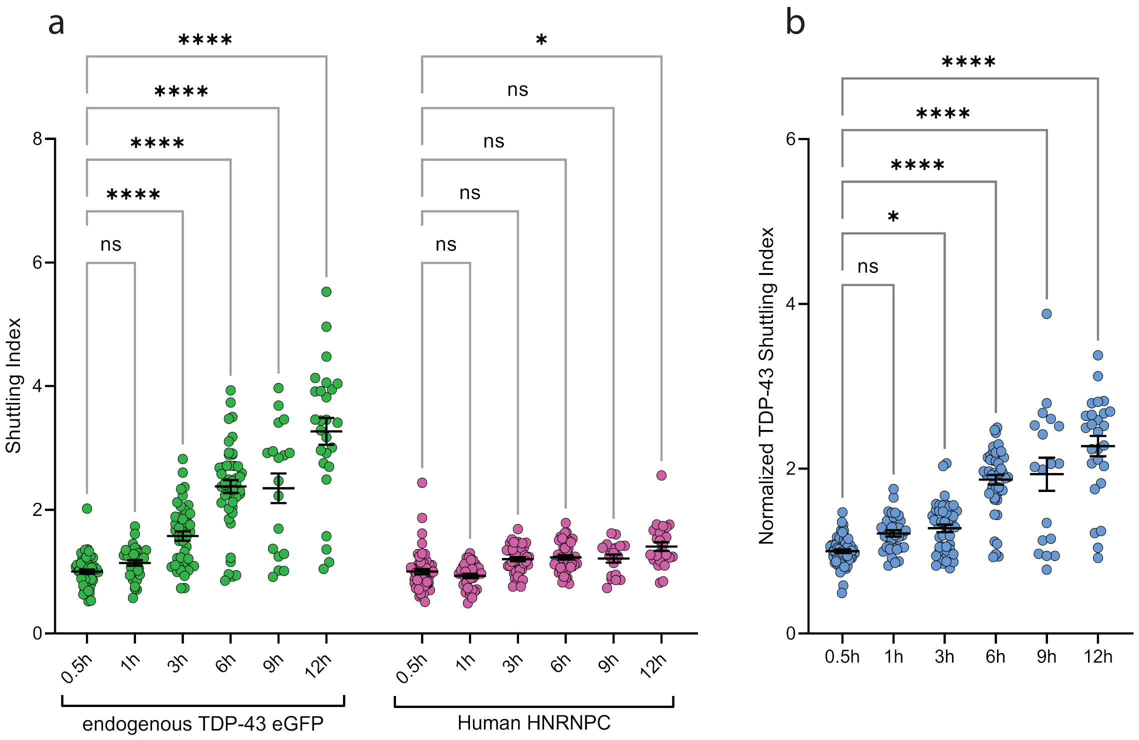


**Supplemental Figure 3: Nuclear export of TDP-43 can be quantified using the heterokaryon assay.**

a) Quantification of the shuttling index of endogenous TDP-43 eGFP (green) shows an increase over time while a known non-shuttling RBP, HNRNPC (pink), shows only minor changes in shuttling rate over time. One-way ANOVA, n = 18-57 heterokaryons, ****p<0.0001, *p<0.5, ns = not significant.

b) Normalizing the raw TDP-43 shuttling index with the HNRNPC shuttling index shows a significant increase over time. One-way ANOVA, n = 18-57 heterokaryons, ****p<0.0001, *p<0.5, ns = not significant.

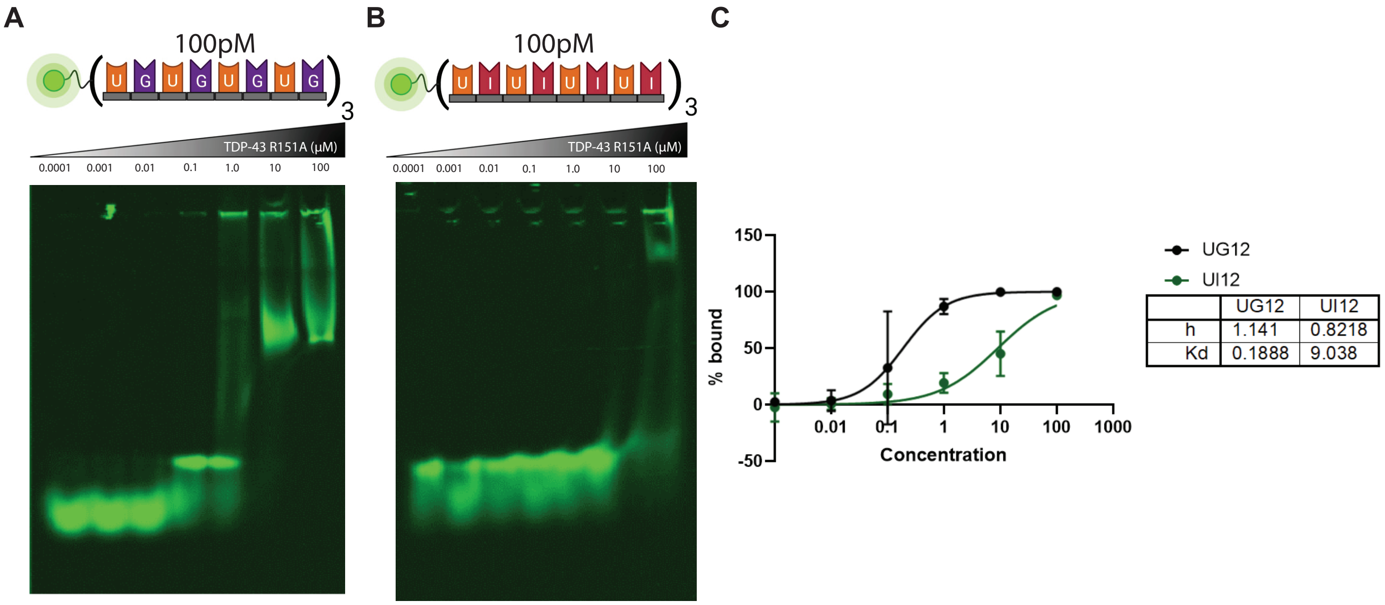


**Supplemental Figure 4: Mutant TDP-43 with altered RNA interactions can still bind UI-repeat RNA oligomers, but not as strongly relative to UG-repeat RNAs.** (A) Increasing concentrations of recombinant TDP-43$^{R151A}$ (0.001 μ M – 100 μM) were mixed with labeled $(UG)_{12}$ (100pM). (B) Increasing concentrations of recombinant TDP-43$^{R151A}$ (0.001 μ M – 100 μM) were mixed with labeled $(UI)_{12}$ (100pM). (C) Kd and h were calculated from 3 different replicates.
